# An equine tendon model for studying intra-tendinous shear in tendons that have more than one muscle contribution

**DOI:** 10.1101/2021.03.19.436206

**Authors:** Nai-Hao Yin, Ian McCarthy, Helen L. Birch

**Affiliations:** Research Department of Orthopaedics and Musculoskeletal Science, University College London, Royal National Orthopaedic Hospital, Brockley Hill, Stanmore HA7 4LP, UK; Pedestrian Accessibility and Movement Environment Laboratory, Department of Civil, Environmental and Geomatic Engineering, University College London, London N19 5UN, UK

**Keywords:** Achilles tendon, Equine AL-DDFT, sub-tendon, biomechanics, interfascicular matrix

## Abstract

Human Achilles tendon is composed of three smaller sub-tendons and exhibits non-uniform internal displacements, which decline with age and after injury, suggesting a potential role in the development of tendinopathies. Studying internal sliding behaviour is therefore important but difficult in human Achilles tendon. Here we propose the equine deep digital flexor tendon (DDFT) and its accessory ligament (AL) as a model to understand the sliding mechanism. The AL-DDFT has a comparable sub-bundle structure, is subjected to high and frequent asymmetric loads and is a natural site of injury similar to human Achilles tendons. Equine AL-DDFT were collected and underwent whole tendon level (n=7) and fascicle level (n=7) quasi-static mechanical testing. Whole tendon level testing was performed by sequentially loading through the proximal AL and subsequently through the proximal DDFT and recording regional strain in the free structures and joined DDFT and AL. Fascicle level testing was performed with focus on the inter-sub-bundle matrix between the two structures at the junction. Our results demonstrate a significant difference in the regional strain between the joined DDFT and AL and a greater transmission of force from the AL to the DDFT than vice versa. These results can be partially explained by the mechanical properties and geometry of the two structures and by differences in the properties of the interfascicular matrices. In conclusion, this tendon model successfully demonstrates that high displacement discrepancy occurs between the two structures and can be used as an easy-access model for study intra-tendinous shear mechanics at the sub-tendon level.

## 1. Introduction

Achilles tendon injury is prevalent in both athletes and the general population [1], but our limited understanding of the pathogenesis has hampered the development of successful prevention and interventional strategies [2, 3]. Injuries are likely initiated from a combination of several factors including the Achilles anatomy [2, 4, 5], the integrity of the extracellular matrix [6, 7] and activity-related external mechanical demands [1, 3]. The susceptibility however relates to the intrinsic Achilles anatomy and the complex force interaction within the tendon bundles [4, 5, 8]. The Achilles tendon has a unique tri-bundled structure formed by two longer and thinner gastrocnemii tendons and a shorter and thicker soleus tendon. Although the different contributing tendons on visual inspection appear to merge into one homogeneous tendon, in fact the tendon fascicles originating from different muscle bellies do not merge or intertwine within the Achilles tendon until reaching the distal calcaneal tuberosity [9-12]. These fascicle groups or ‘bundles’, termed Achilles sub-tendons [13], are believed to facilitate movement efficiency by allowing a certain degree of individual control from contracting muscles to meet the demands from locomotive activities [14-17]. The gastrocnemii sub-tendons occupy the posterior-lateral-anterior region of the Achilles tendon, surrounding the soleus sub-tendon which occupies the medial-posterior region [11, 18]. The gastrocnemii and soleus muscles are activated differently during locomotion [19, 20]. Therefore, the observed mechanical behaviour of the whole Achilles tendon is likely the result of a combination of factors including the morphology and the mechanical properties of sub-tendons, various force level from different muscle bellies, and the interface properties between sub-tendons [8].

Individual control of sub-tendons creates non-uniform displacements within the Achilles tendon, and this non-uniformity has been observed in both passive [21-23] and dynamic [16, 24] movements. Furthermore, this displacement non-uniformity between the sub-tendons shows an age-related and disease-related decline [17, 25, 26]. At the fascicle level, studies on the equine superficial digital flexor tendon (SDFT), an extreme example of an energy-storing tendon, have demonstrated that the high failure strain of this tendon is achieved by having a compliant interfascicular matrix (IFM), which allows sliding of fascicles [27, 28]. This sliding capacity decreases with age due to the stiffening of the IFM [29], presumably imposing premature strains on fascicles and increasing risk of injury. The human Achilles tendon, the most important energy-storing tendon in human locomotion, exhibits a similar fascicle sliding mechanism and compliant IFM [30-32] but whether a significant transmission of force occurs between fascicles remains inconclusive. The matrix between sub-tendons, termed inter-sub-tendon matrix [33], has even more complex requirements than the IFM within a tendon as it is subjected to high asymmetric loads due to the aforementioned individual control from different muscles. The inter-sub-tendon matrix is therefore likely to demonstrate a more extreme specialisation, which may be structural or compositional, than the presumably more homogeneously loaded IFMs within each sub-tendon [33]. The detailed fascicle level mechanical behaviour between human Achilles sub-tendons is difficult to study both *in vivo* and *in vitro* due to the highly complex, individually variable rotatory morphology of the tendon and sub-tendon [12, 18] in addition to the difficulties of obtaining sufficient healthy specimens to study.

Here we have used an equine tendon model for studying the mechanism of intra-tendinous shear between sub-structures – the deep digital flexor tendon (DDFT) and its accessory ligament (AL). The horse is a commonly used and widely accepted animal model for studying human tendon, especially the equine SDFT to represent human Achilles tendon as they have comparable energy-storing functions. However, the SDFT does not possess sub-tendon level structures distal to the metacarpo-phalangeal joint [34] and therefore lacks the ability to provide understanding of shear mechanics between sub-tendons. The equine DDFT and AL have a separate origin and proximal portion but combine to form one, seemingly homogeneous, structure in the more distal part. The DDFT lies just beneath (deep to) the extensively studied SDFT and provides more of a support and positioning role during locomotion compared to the SDFT [35]. The loading pattern differs between the two branches of this structure. The DDFT originates from the deep digital flexor muscle and its loading directly associates with muscular actions; the AL originates from the common palmar ligament of the carpus, so mechanical behaviour is largely dictated by the metacarpo-phalangeal joint angle [35]. Along the descent, the AL gradually flattens and wraps around the DDFT at the mid-metacarpus level. A considerable degree of sliding is required at the junction due to the different mechanical demands and different strain patterns during gait cycles between structures [35]. Distally, the AL completely blends with the DDFT, forming one elliptical adjoined tendon before passing through the metacarpo-phalangeal joint (Fig. 1). Given this bifurcated proximal geometry and the different functional roles, the regional mechanical properties are likely different between the two proximal ends and joined regions.

**Figure 1.**
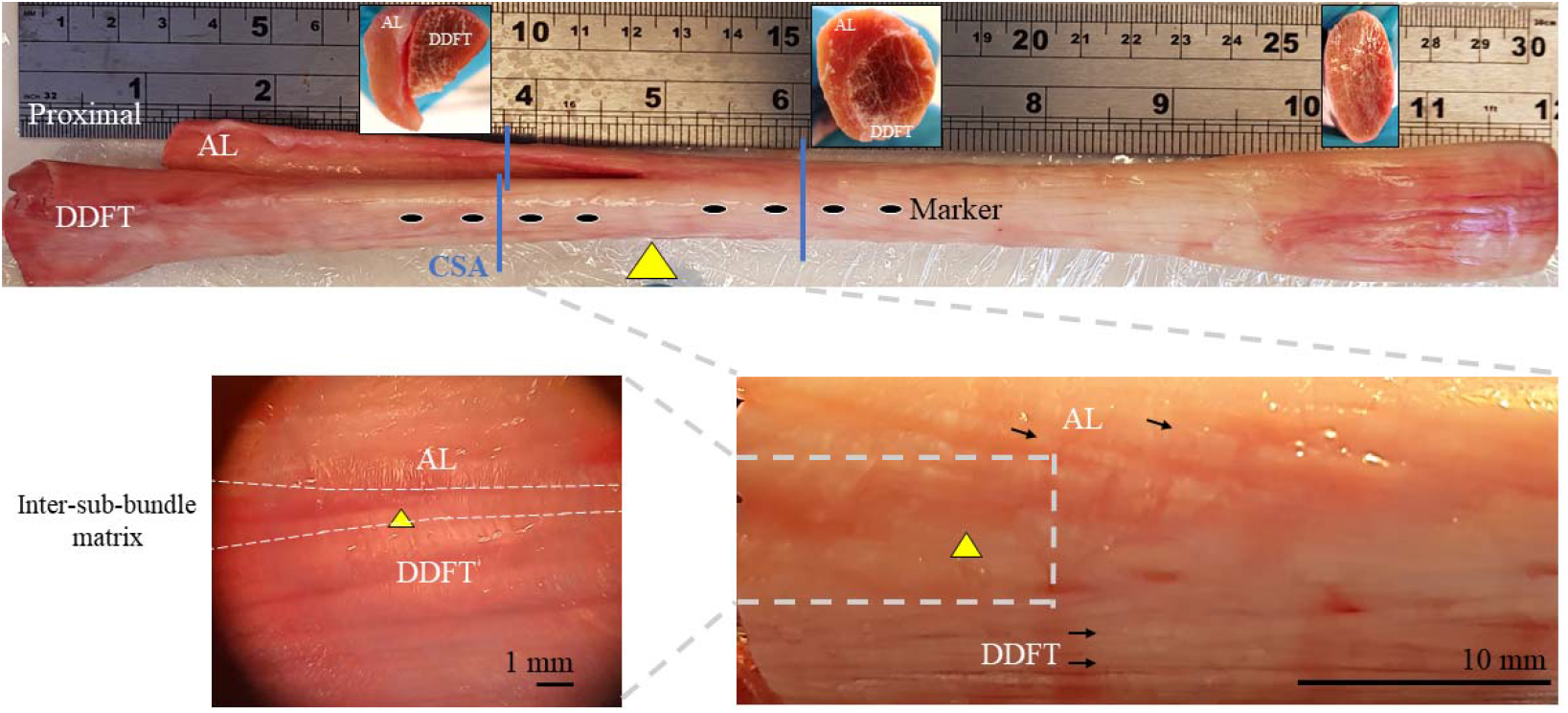
The deep digital flexor tendon (DDFT) and its accessory ligament (AL) and the inter-sub-bundle matrix *(lower left)*. Yellow triangles: the junction between two structures. Blue lines: levels measuring cross-sectional area. Upper row inlets: cross-sections at corresponding longitudinal regions. Black arrows: tendon/ligament fascicles.

It has been demonstrated that the proximal DDFT is a stiffer structure than the proximal AL [35, 36], but no previous study has investigated the mechanical properties at the junction, where the two structures join. The junction is subjected to intermittent high asymmetrical loads transmitted from either the AL or DDFT, resulting in complex force interactions [35]. The shear-related displacement of the less-strained contralateral side will be influenced by the properties of the inter-sub-structure matrix governing the force transmission and the relative sliding behaviour between the two structures [8, 33, 37]. Similarities therefore exist between the human Achilles tendon and the equine DDFT. Both tendons have a sub-tendon/ligament level structure; the difference between tendon and ligament is anatomical rather than a structural or functional feature. Both tendons function to receive and transmit force from different sub-structures and exhibit non-uniform displacements. The requirements of the matrix between Achilles sub-tendons and the matrix between the DDFT and AL are therefore very similar. We propose that studying the AL-DDFT mechanics will improve our knowledge of the sub-tendon level mechanical behaviour within the Achilles tendon.

This study aims to explore the mechanical response at the junction between the AL and DDFT and along the length of the tendon when subjected to asymmetrical loads at both whole tendon and fascicle levels. We hypothesise that the branched nature of this tendinous structure creates high displacement discrepancy at the junction during loading, which is transmitted to the more distal parts of the structure.

## 2. Methods

### 2.1 Sample preparation

In total 17 pairs of equine forelimbs (aged from 3 to 24 years) were collected at a commercial abattoir where horses are euthanised for reasons other than research. The Animal (Scientific Procedures) Act 1986, Schedule 2, does not define collection from these sources as scientific procedures. The DDFT and AL between the carpus and metacarpo-phalangeal joint of each forelimb were harvested within 24 hours and kept frozen at -80°C until further analysis. No sign of tendon pathology was noticed during the harvest. We conveniently chose all the left limbs for biomechanical tests, assuming no significant limb differences between left and right sides.

### 2.2 Whole tendon biomechanical tests

On the test day, specimens (n=10) were thawed at room temperature before measuring the CSA of different regions with the previously described method [38]. The CSA of free-AL and free-DDFT were measured at a point 30 mm proximal to the junction between the AL and the DDFT. The CSA of the joined region was measured at a point 30 mm distal to the junction (Fig. 1).

Non-destructive quasi-static tests were conducted with sequential loading (AL then DDFT) of two proximal free ends. Specimens were vertically mounted, distal part at the bottom and proximal parts at the top, secured by cryoclamps in a screw-driven mechanical testing device (Instron 5967, Instron, MA, US) with a 30 kN load cell. We first mounted the free-AL while keeping the free-DDFT relaxed. After securing the specimen in the device, four markers approximately 25 mm apart were placed directly on the surface along the midline at the regions where the CSAs were measured on the free-AL and free-DDFT and both sides of the joined AL and DDFT (Fig. 1, *black dots, schematic representation*). A preload of 100 N was applied for 1 min, and the effective gauge length was measured between the two freeze lines. The specimen was then preconditioned by applying a 5% strain for 20 cycles using a triangular wave at 1 Hz frequency. After the last cycle, the specimen was returned to the slack position before pulled to 10% strain at a speed of 12 mm/sec. We chose a high-speed loading scenario to represent the *in vivo* physiological loading condition and the influence of viscoelastic behaviour [39] on the mechanical response of the tissue. The free-AL of the specimen was dismounted, and the free-DDFT of the same specimen was then mounted and tested under an identical protocol. We chose to test the AL first as our pilot data showed that the AL is a more compliant structure and therefore the target strain of 10% is reached at a lower load with less chance of damage. Tested specimens were visually inspected after loading through the AL and showed no sign of structural disruption. Subsequently the force-displacement relationship was checked in the data files to ensure no reduction in gradient providing further evidence that the structure had not been damaged.

### 2.3 Calculation

Global force and displacement data were recorded by the mechanical testing machine at 100 Hz. During the test, marker movements were captured at 30 Hz by two video cameras, one facing each side of the specimen. The pixel movements of the four markers (∼25 mm apart) were captured in each video frame and the inter-marker distances were calculated. The regional displacement was then calculated by subtracting the initial inter-marker distances. This regional displacement was then synchronised, using external auditory cues generated by the mechanical testing machine, with force data to plot the regional force-displacement relationship and the linear region was used for calculating stiffness. Regional stress (force / regional CSA) and regional strain (difference in the regional displacement / initial length) were then further calculated and the linear region of the stress-strain relationship was used for calculating Young’s modulus of the free-AL and free-DDFT. An in-house MATLAB (R2020, MathWorks, MA) code was used for video processing, regional displacement measurement, offline synchronisation, and calculations.

### 2.4 Fascicle level mechanical testing

A further seven tendons were used for asymmetric IFM and inter-sub-bundle matrix testing. The peritendinous tissues were carefully removed and the junction of AL and DDFT was visually identified (Fig. 1, *yellow triangles*). For the inter-sub-bundle matrix isolation, a fascicle pair (one fascicle from the AL and one from the DDFT) was visually tracked and separated from proximal (∼5 mm above the junction) to distal, retaining the inter-sub-bundle matrix that binds the AL and DDFT together (Fig. 1, *bottom left*). For the IFM of AL and DDFT, the fascicle pairs were selected at the same junction region of the inter-sub-bundle separation, but with sufficient distance from the inter-sub-bundle matrix (e.g. Fig. 1, *bottom right black arrows*). Fascicle pairs were approximately 30 mm in length and each fascicle was cut leaving a 10 mm interfascicular or inter-sub-bundle matrix for testing [27] (Fig. 2). We could only separate the inter-sub-bundle matrix in the periphery due to the difficultly in tracking fascicle origins at the core. However, previous studies on other equine tendons found no significant difference in mechanical properties between fascicles isolated from the core or the periphery [27].

**Figure 2.**
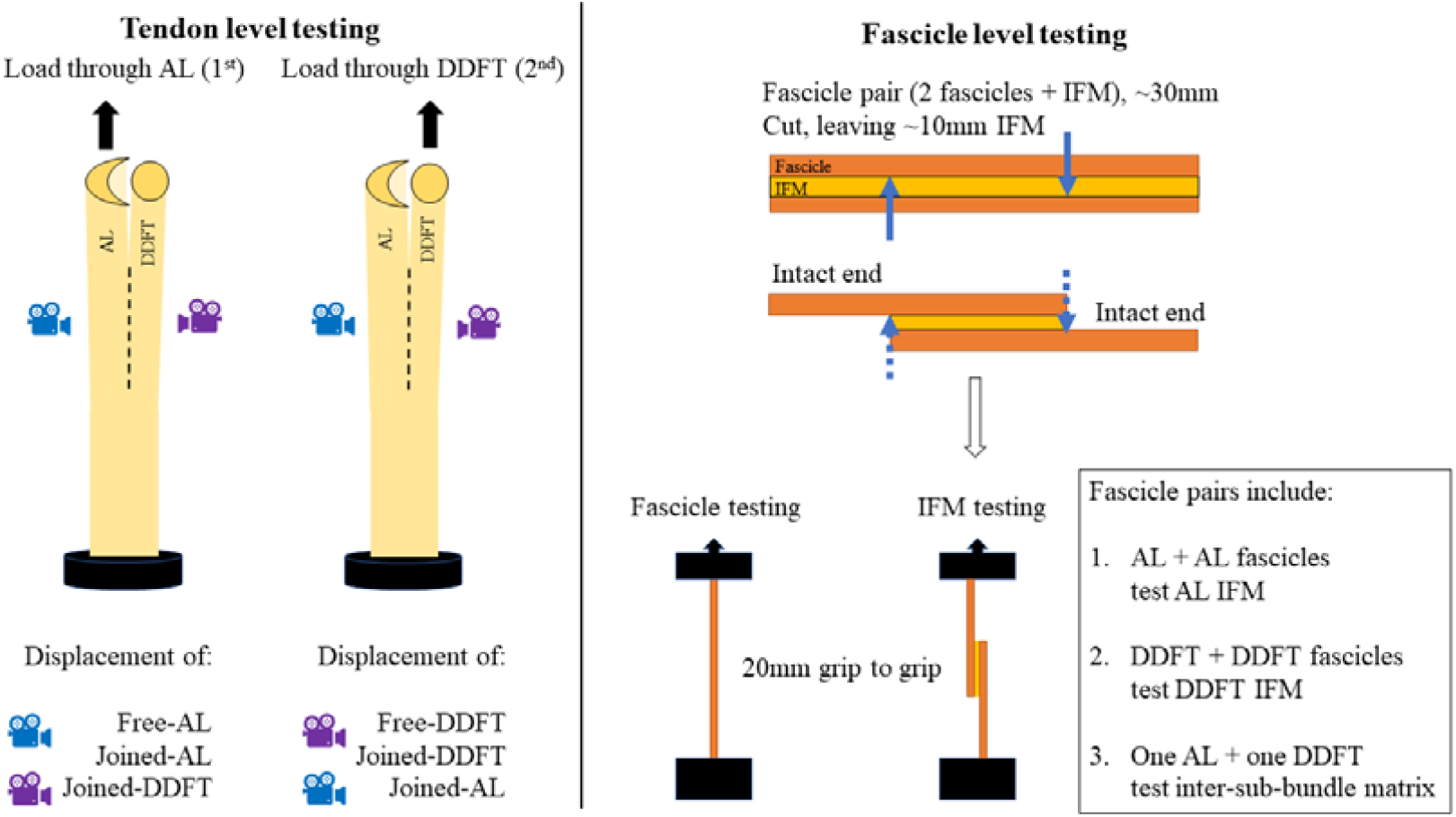
Diagram of tendon level *(left)* and fascicle level *(right)* mechanical testing. The IFM was subjected to shear loading during testing and the fascicles to tensional loading [27, 28].

The intact ends of the fascicles were mounted in a mechanical testing machine (Instron E1000) with a 250 N dynamic load cell (to an accuracy of 0.5% of 1 N and measurement uncertainty less than one-third of the permissible error. 2527-131 Dynacell, Instron.) and custom-made jagged grips. Grip to grip distance was 20 mm (Fig. 2). A preload (0.02 N, as displayed on the mechanical testing machine) was applied and the specimen was preconditioned with 10 cycles of sine waves between 0 to 0.5 mm. The structure was then allowed to slacken before being pulled to failure at 1 mm/s [28]. Force and displacement were recorded at 100 Hz during the test. To normalise data for comparison, the force-displacement relationship was displayed by plotting each 10% of maximal displacement against force. The stiffness was calculated as the slope of the linear region of the force-displacement relationship of each specimen.

Additionally, single fascicles (∼30 mm in length) from AL and DDFT (n=7) were dissected and mechanically tested using an identical testing protocol as the IFM, but with load applied in tension instead of shear (Fig. 2).

### 2.5 Statistical analysis

Statistical analysis was performed using SPSS (v. 26, IBM, NY, US) with two-tailed significant level set at 0.05. A one-way ANOVA with Bonferroni post-hoc adjustment was performed to compare the CSA of three regions. Paired t-tests were performed to compare means of mechanical testing results between free-AL and free-DDFT, between joined-AL and joined-DDFT, and between AL and DDFT fascicles. Strain and stiffness were also compared between free and joined region within the AL or the DDFT. One-way ANOVAs with Bonferroni post-hoc adjustment were conducted for comparing tendon level displacements when loaded in isolation (e.g. free-DDFT, joined-DDFT, and joined-AL when DDFT was loaded) and also interfascicular mechanical properties (AL IFM, DDFT IFM, and inter-sub-bundle matrix).

## 3. Results

### 3.1 Tendon level mechanical properties

The CSA (n=10) of the joined region (159±30 mm^2^) was significantly larger than the free ends of both AL (96±32 mm^2^) and DDFT (89±28 mm^2^) but smaller than the combined CSAs. The mechanical testing results of the tendon level (n=7) and fascicle level (n=7) are summarised in Tables 1-3. The mechanical testing data from 3 limbs were discarded due to poor image quality or clamp slippage during the whole tendon testing.

**Table 1.** Mechanical testing results when load was applied through the AL

|  | Global | Regional |  |  |
| --- | --- | --- | --- | --- |
|  |  | Free-AL | Joined-AL | Joined-DDFT |
| Maximal force (N) | 4320±1684 |  |  |  |
| Displacement (mm) | 13±1.8 | 5.0±1.4<sup>a, b</sup> | 3.6±1.5<sup>b</sup> | 1.9±0.8 |
| Stress (MPa) |  | 44.8±15.4<sup>c</sup> |  |  |
| Strain (%) | 10±0.0 | 6.8±2.0<sup>a, b, c</sup> | 4.0±1.3<sup>b</sup> | 1.9±0.8 |
| Young's modulus (MPa) |  | 639±244<sup>c</sup> |  |  |
| Stiffness (N/mm) |  | 841±327<sup>c</sup> | 1247±836 |  |

Data are shown as mean ± SD (n=7). a: Significant difference from joined-AL. b: Significant difference from joined-DDFT. c: Significant difference from the same location as DDFT (Table 2).

When the AL was loaded, the machine recorded lower maximal force (global) compared to the DDFT loading condition. The regional displacement and strain (measured by cameras) showed significant regional differences between free-AL, joined-AL, and joined DDFT, which receives load only through shear. The stiffness of the free-AL was significantly lower than the joined-AL when the AL was directly loaded (Table 1).

When the DDFT was loaded in isolation, the displacement, strain, and stiffness were similar between the free- and joined-DDFT, contrary to the AL loading condition (Table 2). Comparing the two loading conditions revealed that the free-DDFT was a stiffer structure than the free-AL and had a higher material stiffness (Fig. 3). Interestingly, when comparing the displacement of the contralateral joined regions, the force transmittance pattern was different between AL and DDFT loading conditions. When the DDFT was directly loaded, the joined-AL remained stationary and started to elongate only after a substantial force was imposed on the DDFT (Fig. 4, *right*). When the AL was directly loaded, the joined-DDFT displaced proportionally to the joined-AL strain (Fig. 4, *left*). Maximal displacement discrepancy at the junction was approximately 3.0 mm (DDFT loaded, 3920 N) and 1.7 mm (AL loaded, 4192 N).

**Table 2.** Mechanical testing results when load was applied through the DDFT.

|  | Global | Regional |  |  |
| --- | --- | --- | --- | --- |
|  |  | Free-DDFT | Joined-DDFT | Joined-AL |
| Maximal force (N) | 5370±2108 |  |  |  |
| Displacement (mm) | 14±2.4 | 4.1±1.4<sup>b</sup> | 4.2±1.1<sup>b</sup> | 1.2±1.5 |
| Stress (MPa) |  | 60.7±26.9<sup>c</sup> |  |  |
| Strain (%) | 10±0.0 | 5.3±1.5<sup>b, c</sup> | 4.9±1.8<sup>b</sup> | 1.4±1.8 |
| Young's modulus (MPa) |  | 1158±459<sup>c</sup> |  |  |
| Stiffness (N/mm) |  | 1387±639<sup>c</sup> | 1224±486 |  |

Data are shown as mean ± SD (n=7). a: Significant difference from joined-DDFT. b:
Significant difference from joined-AL. c: Significant difference from the same location as AL
(Table 1).

**Figure 3.**
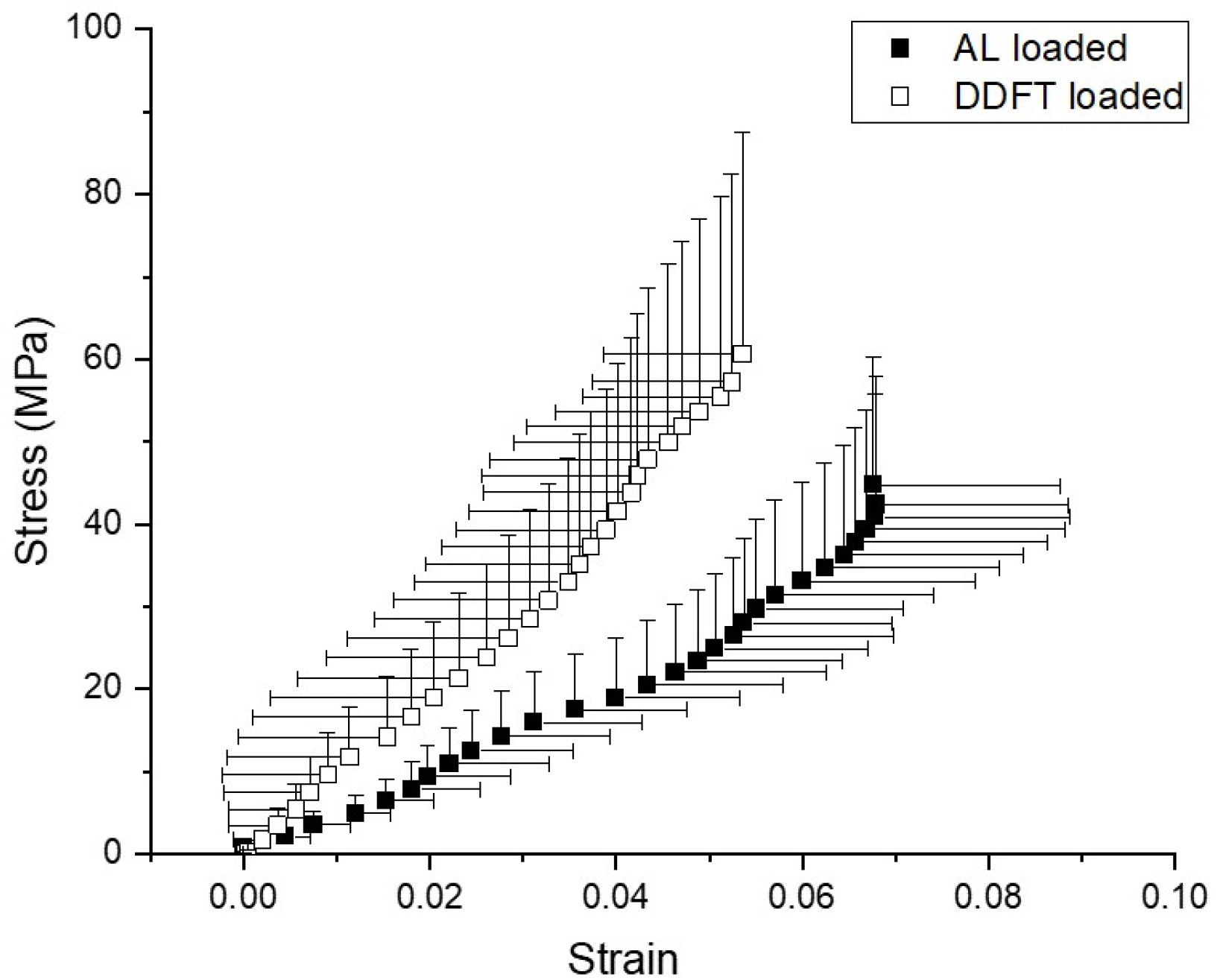
Averaged (n=7) stress-strain relationship of free-AL and free-DDFT. Error bars represent standard deviation and shown at one side for clear visualisation.

**Figure 4.**
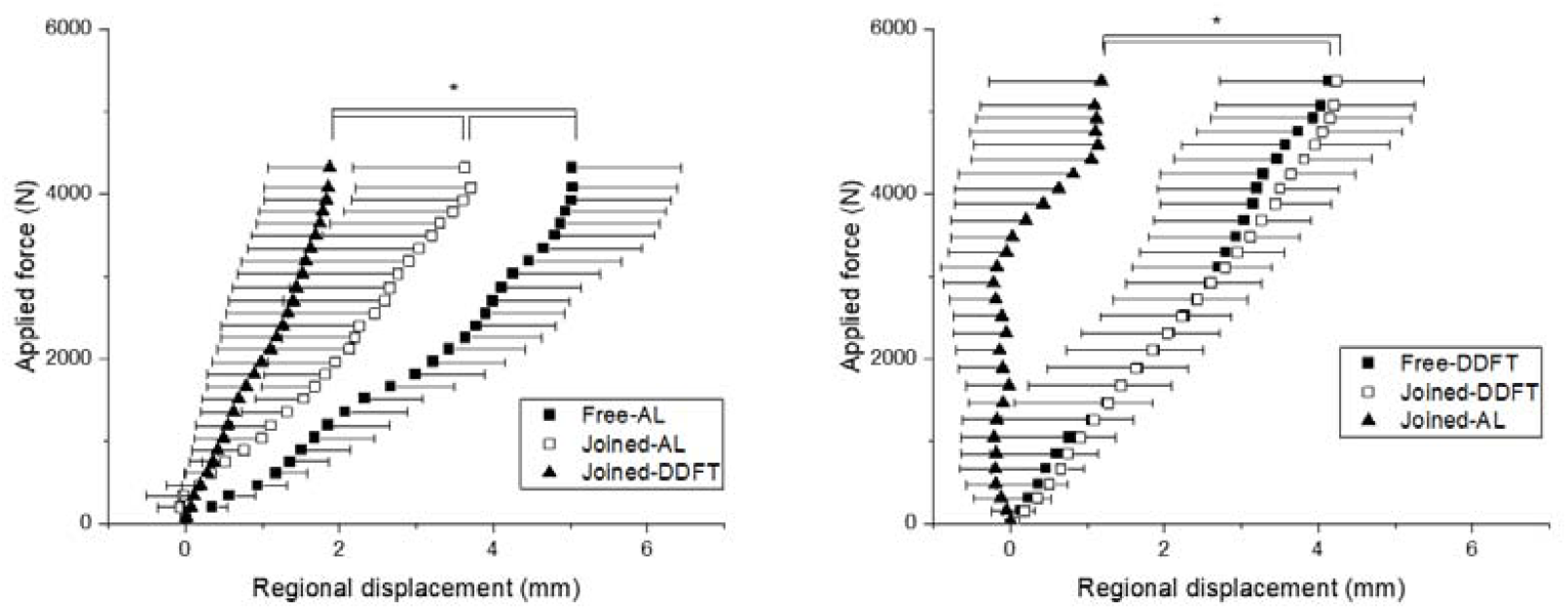
Averaged (n=7) force-displacement relationship when AL *(left)* and DDFT *(right)* were loaded in isolation. The contralateral side (joined-DDFT when AL loaded, joined-AL when DDFT loaded; *closed triangles*) was shear loaded alone. Asterisks: significant difference in maximal displacement. Error bars represent standard deviation of regional displacement and are shown at one side for clear visualisation.

### 3.2 Fascicle level mechanical properties

AL and DDFT fascicles exhibited similar failure force and maximal displacement (Table 3) and their mechanical properties were not significantly different. The averaged AL IFM appeared to have a slightly higher failure force and stiffness than the DDFT IFM and the inter-sub-bundle matrix, but the differences were not statistically significant (Table 3). The relationship between load and displacement however differs significantly; at 20% and 30% of normalised displacement levels, AL IFM exhibited significantly higher force than the inter-sub-bundle matrix (Fig. 5). At higher normalised displacement levels differences were not significant.

**Table 3.** Mechanical properties of fascicles and Interfascicular matrices

|  | Interfascicular matrices |  |  | Fascicles |  |
| --- | --- | --- | --- | --- | --- |
|  | AL | DDFT | Inter-sub-bundle | AL | DDFT |
| Failure force (N) | 2.2±1.1 | 1.4±0.8 | 1.1±0.8 | 5.3±0.9 | 4.9±2.5 |
| Failure strain (%) |  |  |  | 7.5±5.5 | 7.5±4.5 |
| Maximal displacement (mm) | 1.9±0.7 | 1.7±0.5 | 1.8±0.7 | 1.5±1.1 | 1.5±0.9 |
| Stiffness (N/strain%) |  |  |  | 0.7±0.2 | 0.7±0.6 |
| Stiffness (N/mm) | 1.6±0.7 | 1.3±0.7 | 0.9±0.6 | 4.3±2.3 | 3.2±2.2 |

Data are shown as mean ± SD (n=7). N.B displacement for interfascicular matrix is given in mm due to un-defined effective gauge length.

**Figure 5.**
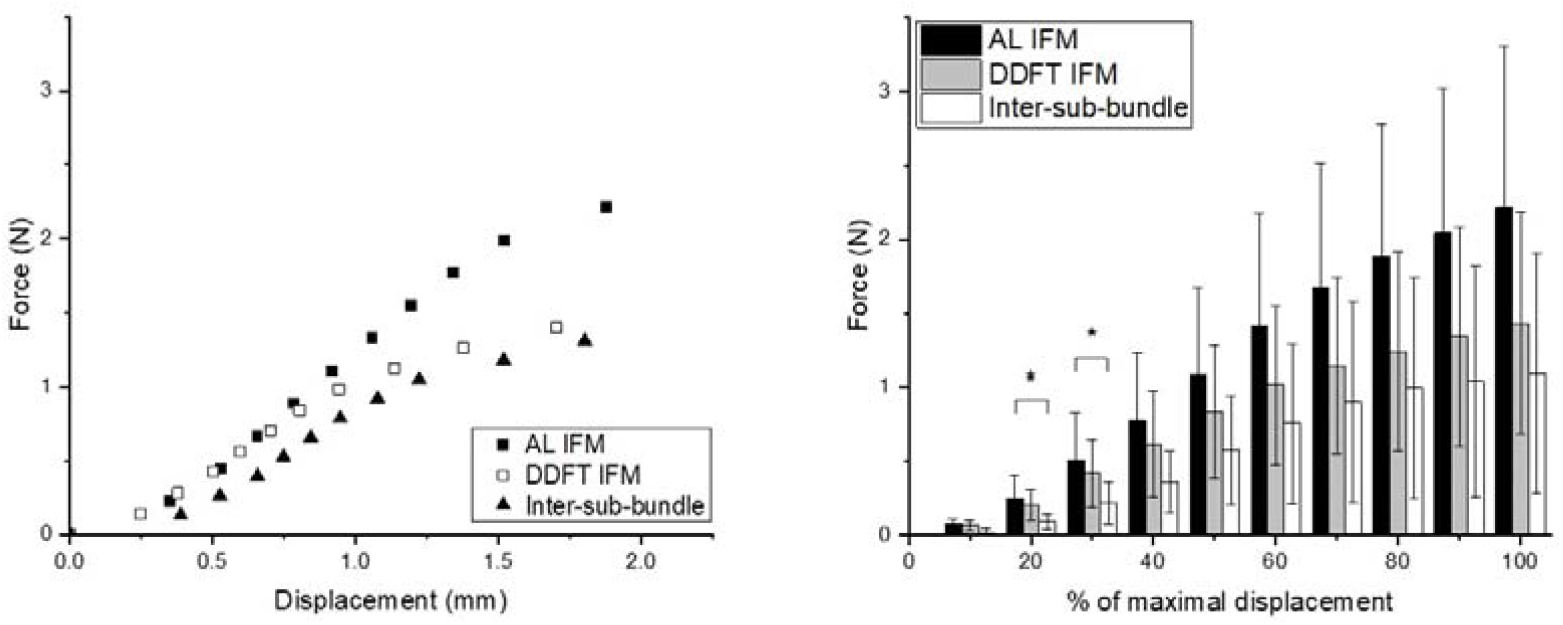
Relationship between force and displacement *(left)* and between force and normalised displacement *(right)* of interfascicular matrices (n=7). Error bars are not shown in the force-displacement curves *(left)* for better visual presentation of the data. Asterisks: significant difference after post-hoc analysis (*p*<0.017).

## 4. Discussion

Our results demonstrate that this equine tendon model exhibits large displacement discrepancy at the junction when subjected to asymmetry loads from different branches as expected. Interestingly, the results confirm our hypothesis and demonstrate that this discrepancy is maintained in the more distal parts of the structure where the AL and DDFT appear to be tightly bound. Since similar structural and functional features exist between the human Achilles tendon and this equine tendon model, exploiting this new tendon model may help us understand the mechanisms behind the complex mechanical behaviour at the sub-tendon level [30].

The mechanical properties obtained from the free-AL and free-DDFT are comparable with previous studies on isolated AL [40] and DDFT [35] and, similar to previous reports, demonstrated high individual variability at the tendon level [29, 40] and fascicle level [28, 29], reflecting the nature variation between animals. The higher material stiffness of the DDFT fulfils its mechanical role of positioning distal joints in flexion during galloping, a function similar to the equine digital extensor tendons and the human tibialis anterior tendon. The differences in stiffness between the free-DDFT and the free-AL was not reflected at the fascicle level, at least at the joined region. Future research is warranted to elucidate the origin of the difference in mechanical properties of the AL and DDFT, such as the inter-structure matrix composition or the ratio between IFM and fascicles. Interestingly, in our study the palmar and dorsal aspects of the joined AL and DDFT structure behaved differently compared to the proximal free regions. The regional strain of the joined-AL was significantly lower than the free-AL, suggesting a large proportion of force transmitted through shear-loaded alone to the contralateral joined-DDFT, the regional strain of which was lower but closely related to the joined-AL. On the contrary, the regional strain of free- and joined-DDFT were almost identical when loaded in isolation, suggesting a more homogeneous structure throughout the DDFT and a lower force transmitting capacity at the junction compared to force transmission from the AL to the DDFT. The adjacent joined-AL displaced surprisingly little even when the DDFT was subjected to a substantial level of force (∼4000 N).

This force transmission discrepancy at the junction between different loading conditions can be partially explained by the mechanical properties of DDFT and AL. To a first approximation they can be considered as springs in parallel. However, in a classical parallel spring system, the displacements of joined-DDFT and joined-AL would be the same. Another explanation of this unique behaviour at the junction is the different geometry between the AL and DDFT. The AL gradually flattens and wraps around the DDFT when descending, forming a C-shape structure surrounding the O-shape DDFT at the junction. (Fig. 1, *upper row inlets*). Due to Poisson’s effect, the C-shape AL when loaded could exert compressive force on the DDFT, transmitting force through the inter-sub-bundle matrix resulting in displacement of the joined-DDFT. When load is applied through the DDFT however, the decreasing CSA of the O-shape DDFT when loaded may make it less likely to exert force on the joined-AL. Although this theory alone does not explain why the joined-AL eventually displaced when high loads were imposed on the DDFT. Understanding how the mechanical properties and geometry of sub-components within one tendon structure affects mechanical behaviour will be a breakthrough in understanding Achilles tendon mechanics since a similar mechanical property differences and geometry feature (C-shape gastrocnemii surrounds the soleus sub-tendon) exists in Achilles sub-tendons [8, 10-12]. A third explanation is that the orientation of load bearing fibres in the inter-sub-bundle matrix is such that they come under tension much later when loaded through the DDFT than when loaded through the AL resulting in a difference in timing of force transmission between the two structures.

Fourthly, the relative mechanical properties of the interfascicular and inter-bundle matrix could also contribute to this unique behaviour seen at the junction. Given the large differences in the force-displacement relationship between the AL IFM and inter-sub-bundle matrix, a near-failure force for the inter-sub-bundle matrix could only result in approximately half of the displacement in the AL IFM. On the contrary, since the force-displacement relationships between DDFT IFM and inter-sub-bundle matrix were similar and both were more compliant and weaker than the AL IFM, a fully loaded AL could result in a substantial displacement of the DDFT, showing a more synchronised displacement between the shear-loaded joined-DDFT and the directly loaded joined-AL. However, these inter-fascicular differences in mechanical properties were only statistically significant at initial displacements (<30%). This may be due to the small sample size, mixed ages, and the inherent variation between individual animals. We were not able to measure the thickness or CSA of the inter-fascicular matrices or to maintain identical IFM quantity (volume) during our test. Although we did not find significant correlation between age and measured tendon and fascicle level mechanical properties for both AL and DDFT, ageing has been proposed to have a significant influence on the IFM tensile mechanical properties [27, 28] and fascicle and fibre level viscosity [41]. A greater adhesion of inter-sub-bundle matrix would drastically affect force transmission at the AL-DDFT junction. Future studies may benefit from obtaining a larger sample size with a wide age range to detect potential fascicle IFM and inter-sub-bundle matrix level differences. Our measured fascicle level mechanical properties were similar to those reported for the SDFT [27, 28], and these results also imply that interfascicular differences have a greater influence on whole structure mechanical properties than the fascicle differences. Additional research is warranted to explore whether fascicle sliding also plays an important role in the DDFT during loading as there are important differences in function and composition between the SDFT and DDFT [35, 42]. We could only confidently dissect out fascicles at the periphery due to the difficulty in tracing the fascicular origin at the core. Previous studies however suggest that the fascicle and IFM mechanical properties are similar between at the core and the periphery in other equine tendons [27]. We only reported the force-displacement relationship since we were unable to accurately measure the CSA of the tested fascicles. The actual material strength of fascicles is supposedly greater than the IFM [31]. Future study could also investigate the influence from the lower hierarchies (fibril or fibre level) on the fascicle and sub-tendon level mechanical behaviours [39, 43-46].

In some specimens, an initial negative displacement of the joined-AL was observed when the DDFT was loaded. This apparent negative displacement could result from an error from our image-based measurement. The highest negative displacement measured under 4000 N was - 0.017 mm (maximal positive displacement: 0.022 mm). We believe this potential measurement error did not significantly affect the reported outcome. Alternatively, the negative displacement may result from a bending of the structure when load is initially applied to the contralateral side. It is also worth noting that macroscale strain does not fully represent the microscale strain [47]. Therefore, our measured external regional strain may not fully represent the internal strain of the highly heterogenous tendon structure.

Our mechanical testing results can give an insight into why injury to the AL is more prevalent than injury to the DDFT [48, 49]. During galloping, the AL is taut rapidly when the carpal joint moves from flexion to extension while the DDFT remains taut continuously. Taken together that the AL is less stiff than the DDFT and the free-AL is more compliant than the joined-AL, the rapid and high-intensity loading imposed on the whole heterogeneous AL structure may produce localised high strains proximal to the junction and high shear strain at the junction. Indeed, injury to the AL proper (free-AL region) is more frequently reported than injuries to the free-DDFT or below the junction level [48]. It is also worth noting that adhesion between AL and DDFT is sometimes reported with AL desmitis [48]. Adhesion at the junction may be a reaction to injury and may interrupt the unique mechanical response in this tendon complex, but the exact consequence is unknown.

The human Achilles tendon is also likely to receive considerable shear forces and it has been suggested that high shear between Achilles sub-tendons is a risk factor for developing tendinopathy ([50-52] and review [53]). The soleus muscle-tendon unit constantly activates during bipedal movements. It is composed of mainly type I fibres and has a high fatigue resistance to allow continuous support of body weight during stance and walking. Gastrocnemii have shorter muscle bellies and longer tendinous parts compared to the soleus and are involved during explosive movements such as jumping or at the push-off phase during late stance [19, 20]. Moreover, the length of gastrocnemii is affected by both knee and ankle joint angle while the length of soleus, a one-joint muscle, is dictated only by the ankle joint angle. These anatomic and functional differences of human triceps surae muscles can create large displacement discrepancy within the Achilles tendon, with the highest potential difference between the soleus and gastrocnemii sub-tendons. Younger Achilles tendons exhibit more non-uniformity in sub-tendon displacements compared to aged tendons [15, 16]; therefore, shear strains at the interfaces should be greater in younger than aged Achilles tendons. In support, one study suggested that longitudinal tears within the Achilles tendon are an under-reported pathology prevalent in young, elite-level athletes [51]. Elite athletes are also subjected to high training doses and participate in competitive sports, both of which are likely to increase the chance of asymmetrical loadings by either rapidly changing knee and ankle positions or developing training-induced fatigue to the gastrocnemii. Our proposed equine tendon model can help researchers to systematically study the shear-related mechanical responses at the interfaces between structures and understand how tendons adapt their extracellular matrix composition and organisation to withstand this high internal displacement discrepancy. Understanding these responses will facilitate the development of novel prevention and treatment strategies targeting Achilles tendon injuries.

## 5. Conclusion

The AL-DDFT tendon complex shows distinct structural and regional mechanical properties and exhibits large displacement discrepancy at and below the junction when subjecting to asymmetrical loaded through difference branches. This tendon model can be studied macroscopically using conventional mechanical testing devices at both whole tendon and fascicle level, providing tendon researchers an easy-access model to study intra-tendinous shear mechanics, such as the shear between human soleus and gastrocnemii sub-tendons. The relatively low injury rate of the DDFT compared to the SDFT, despite having high internal displacement discrepancy, may suggest a specialised adaptation strategy exists at the junction. Studying the mechanism behind these shear-related mechanical responses and the adaptation strategy can improve our understanding of human Achilles tendon which, in a manner similar to the AL-DDFT, is constantly subjected to asymmetrical loads from different origins and exhibits non-uniform displacements during activities.

## Acknowledgements

N.Y. thanks the generosity of the Taiwanese government in providing a scholarship for his PhD studies.

## Abbreviations

AL: the accessory ligament (of the deep digital flexor tendon)
CSA: cross-sectional area
DDFT: deep digital flexor tendon
IFM: interfascicular matrix
SDFT: superficial digital flexor tendon

